# 661W photoreceptor cell line as a cell model for studying retinal ciliopathies

**DOI:** 10.1101/479212

**Authors:** Gabrielle Wheway, Liliya Nazlamova, Dann Turner, Stephen Cross

## Abstract

The retina contains several ciliated cell types, including the retinal pigment epithelium (RPE) and photoreceptor cells. The photoreceptor cilium is one of the most highly modified sensory cilia in the human body. The outer segment of the photoreceptor is a highly elaborate primary cilium, containing stacks or folds of membrane where the photopigment molecules are located. Perhaps unsurprisingly, defects in cilia often lead to retinal phenotypes, either as part of syndromic conditions involving other organs, or in isolation in the so-called retinal ciliopathies.

The study of retinal ciliopathies has been limited by a lack of retinal cell lines. RPE1 retinal pigment epithelial cell line is commonly used in such studies, but the existence of a photoreceptor cell line has largely been neglected in the retinal ciliopathy field. 661W cone photoreceptor cells, derived from mouse, have been widely used as a model for studying macular degeneration, but not described as a model for studying retinal ciliopathies such as retinitis pigmentosa.

Here, we characterise the 661W cell line as a model for studying retinal ciliopathies. We fully characterise the expression profile of these cells over many passages, using whole transcriptome RNA sequencing, and provide this data on Gene Expression Omnibus (GEO) for the advantage of the scientific community. We show that these cells robustly express the majority of markers of cone cell origin, including short wave and medium wave opsin. Western blotting confirms expression of selected markers.

Using immunostaining and confocal microscopy, alongside scanning electron microscopy, we show that these cells grow long primary cilia, reminiscent of photoreceptor outer segments, and localise many cilium proteins to the axoneme, membrane and transition zone. Immunostaining shows that opsins are localised to the base of this primary cilium. We show that siRNA knockdown of cilia genes Ift88 results in loss of cilia, and that this can be assayed by high-throughput screening. We present evidence that the 661W cell line is a useful cell model for studying retinal ciliopathies.

## Introduction

The sensory primary cilium is an important non-motile cellular organelle, responsible for detecting changes in the extracellular environment and transducing signals to allow the cell to respond accordingly. The retina contains several ciliated cell types, including the retinal pigment epithelium (RPE) and photoreceptor cells. There are two types of photoreceptors; rods and cones, which differ in their shape and the photopigment they contain. Both cell types contain an inner segment, where the nuclei and other organelles are located. Extending from the apical surface of this inner segment is the connecting cilium, which contains a proximal region analogous to the transition zone of primary cilia on other cell types, and a distal region which is unique to the photoreceptor connecting cilia (Dharmat et al., 2018). At the end of the connecting cilium is a huge elaboration of stacks of membrane where the photopigment molecules are located; termed the outer segment (Sjöstrand, 1953). Rods have a long, thin rod-like outer segment containing rhodopsin (Wheway et al., 2014), and cones have a shorter conical outer segment containing opsins which absorb different wavelengths to allow colour vision (Figure 1). Cone outer segments are often described as having folds of membrane continuous with the plasma membrane rather than disks separate from the plasma membrane but this is only true in lower vertebrates (Pearring et al., 2013, May-Simera et al., 2017).

**Figure 1.**
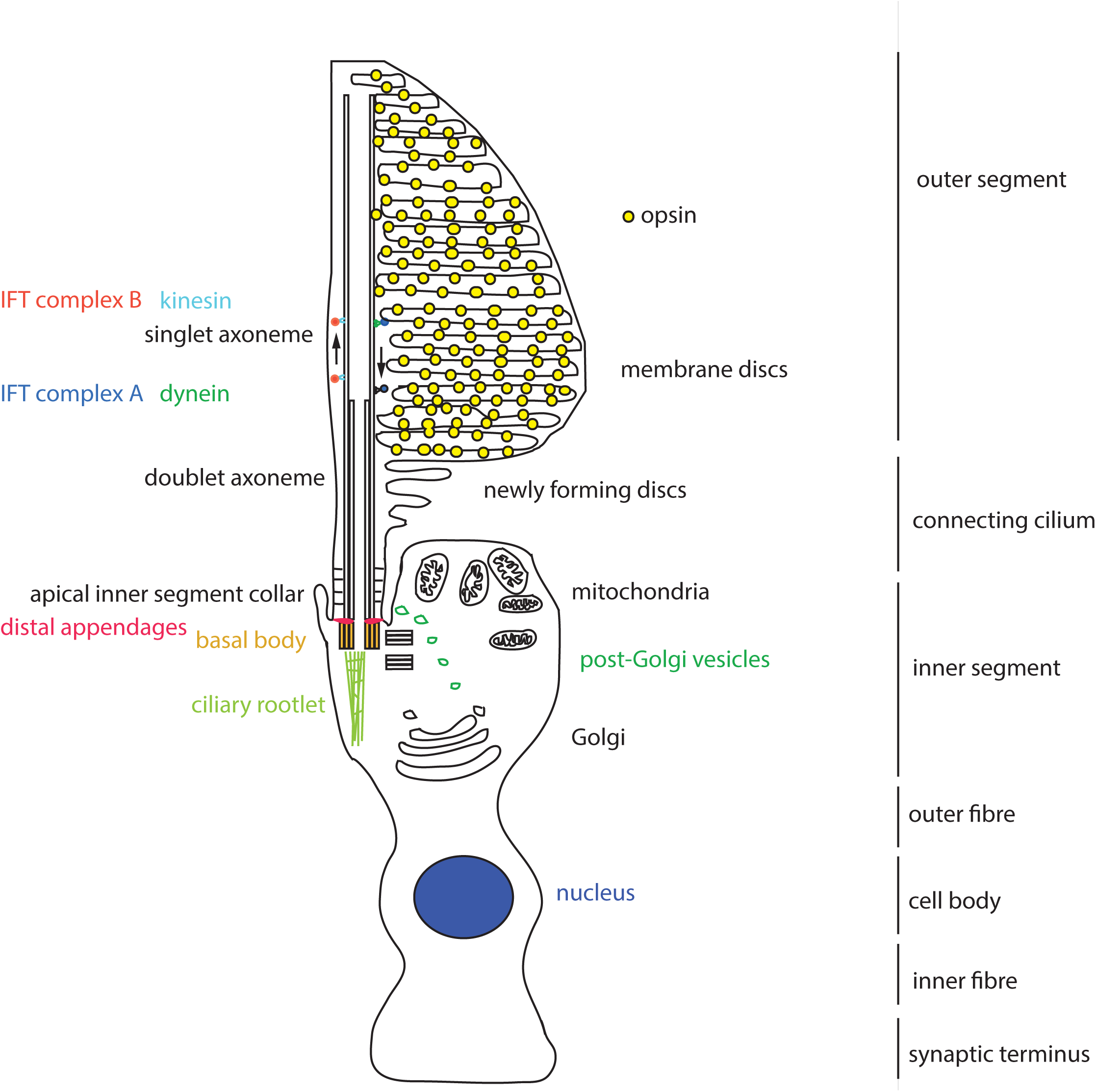
Schematic representation of a cone photoreceptor cell and localization of ciliary proteins. The schematic represents the cone photoreceptor cell outer segment, connecting cilium, inner segment, outer fiber, cell body, inner fiber and synaptic terminus. A number of key components of the ciliary apparatus are colour coded and indicated. The IFT complex A (blue) and complex B (red) are represented in the magnified inset.

Proteins are moved from the site of production, in the inner segment, to the site of light absorption, in the outer segment, along the connecting cilium via a process known as intraflagellar transport (IFT) (Ishakawa and Marshall, 2017). The connecting cilium consists of an axoneme of nine microtubule doublets nucleated at the base by a triplet microtubule structure named the basal body. This structure is derived from the mother centriole, at the apical surface of the inner segment. The axoneme extends into the outer segment, converting to singlet microtubules toward the distal end, often reaching near the distal tip of the cone outer segment and at least half-way along the rod outer segment (Roof et al., 1991). The proximal region of the axoneme is stabilized by posttranslational modifications such as glutamylation and acetylation, and is turned over at the distal end as membranes are replaced at the distal end of the outer segment, particularly in cones (Eckmiller, 1996).

Collectively, the connecting cilium and outer segment are termed the photoreceptor cilium, and this is the most highly modified and specialised sensory cilium in the human body (Wheway, et al., 2014).

Perhaps unsurprisingly, defects in cilia often lead to retinal phenotypes, either as part of syndromic conditions involving other organs, or in isolation in the so-called retinal ciliopathies (Bujakowska et al, 2017). Non-syndromic retinal ciliopathies include several genetic subtypes of retinitis pigmentosa (RP), Leber Congenital Amaurosis (LCA) and conerod dystrophy (CORD).

RP is the most common cause of inherited blindness, affecting up to 1:2000 people worldwide (Golovleva et al 2010; Sharon and Banin, 2015), and is characterised by night blindness and loss of peripheral vision due to degeneration of rod photoreceptor cells, often progressing to loss of central high acuity vision as cone receptors are also affected (Verbakel et al., 2018). It is normally diagnosed in the third or fourth decade of life, although age of onset and severity vary widely. It can occur in isolation, or as part of syndromes such as Usher syndrome, Bardet Biedl syndrome and Joubert syndrome. The condition is extremely genetically heterogeneous, with 64 genes identified as causes of non-syndromic RP, and more than 50 genes associated with syndromic RP (RetNet https://sph.uth.edu/retnet/sum-dis.htm) and can be inherited in an autosomal dominant, autosomal recessive, or X-linked manner.

Of the genetic causes of X-linked non-syndromic RP, OFD1, RP2 and RPGR encode cilia proteins. Within autosomal dominant non-syndromic RP (ADRP), RP1 and TOPORS (RP31) encode known cilia proteins. At least 13 genetic causes of autosomal recessive non-syndromic RP (ARRP) encode cilia proteins, including FAM161A (RP28), TTC8 (RP51), C2orf71 (RP54), ARL6 (RP55), MAK (RP62), NEK2 (RP67), BBS2 (RP74), IFT140 (RP80), ARL2BP, RP1L1, C8orf37, CC2D2A and IFT172.

LCA is the most common genetic cause of childhood blindness, with an estimated worldwide incidence between 1:33,000 (Alstrom 1957) and 1:81,000 live births (Stone 2007). It accounts for 20% of eye disease in children attending schools for the blind (Schappertkimmijser and Vandenbosch 1959). Patients with LCA are born with severe visual impairment which is normally diagnosed within a few months of birth by greatly reduced or non-recordable electroretinogram results. Alongside poor vision are nystagmus (involuntary movement of the eyes) and slow to no pupillary response. Although born with already poor eyesight, some patients with LCA undergo further deterioration of vision in adult life, with retinal pigmentary changes often occurring later in life (Heher et al. 1992). It can occur in isolation, or as part of syndromes such as Senior-Loken syndrome. LCA is also genetically heterogeneous, with 13 known genes associated with autosomal recessive LCA, and one gene associated with autosomal dominant LCA (RetNet https://sph.uth.edu/retnet/sum-dis.htm).

Five genetic subtypes of LCA are known retinal ciliopathies. *LCA5* encodes lebercilin, a ciliary transport protein (den Hollander et al. 2007), *LCA6* encodes RPGRIP1, a ciliary transition zone protein (Dryja et al. 2001), *LCA10* encodes CEP290, a transition zone protein which is also mutated in numerous syndromic ciliopathies (den Hollander et al., 2006) and *LCA15* encodes IQCB1/NPHP5 which interacts with CEP290, localises to the transition zone and is required for outer segment formation (Estrada-Cuzcano et al., 2010; Ronquillo et al., 2016). All of these proteins localise to the connecting cilium of photoreceptor cells. CLUAP1 (IFT38) is also a cause of LCA (Soens et al., 2016), and plays a central role in photoreceptor ciliogenesis (Lee et al., 2014).

Cone-rod dystrophies (CRD) are rare degenerative conditions with an estimated incidence of 1:40,000 (Hamel et al., 2000). The condition is characterised by loss of cone photoreceptors, leading to loss of central, high acuity vision, disruption of colour vision (dyschromatopsia) and photophobia, sometimes followed by degeneration of rod photoreceptors, causing night blindness and tunnel vision. It is normally diagnosed in the first decade of life (Hamel, 2007). It can occur as an isolated condition or as part of the syndromic ciliopathy Alström syndrome (Collin et al., 2002; Hearn et al., 2002). CRDs are also genetically heterogeneous, with 16 autosomal recessive and 5 autosomal dominant genes having been identified as causing CRD (https://sph.uth.edu/retnet/sum-dis.htm). Of these, at least seven encode cilia proteins (RAB28 (CORD18), C8orf37 (CORD16), CEP78, POC1B, IFT81, RPGRIP1 and TTLL5).

In total, at least 30 cilia genes have been identified as genetic causes of non-syndromic retinal dystrophies, and this number continues to grow. New ciliary causes of retinal dystrophies continue to be discovered, and new links are made between cilia and retinal conditions not previously considered to be retinal ciliopathies. For example, a recent whole genome siRNA knockdown screen in a ciliated cell line identified PRPF6, PRPF8 and PRPF31, known causes of RP, as cilia proteins (Wheway et al., 2015), offering new perspectives on a poorly-understood form of RP.

Clearly, the cilium is of central importance to retinal development and function, with defects in large numbers of cilia proteins leading to various inherited retinal dystrophies. Retinal dystrophies remain extremely difficult to treat, with very few, if any, treatment options for the vast majority of patients, with the exception of RPE65, CEP290 and GUY2D gene therapy in LCA (DiCarlo et al., 2018). In order for this situation to improve, better understanding of the cell biology and molecular genetics of retinal dystrophies, including retinal ciliopathies, is required.

This requires robust, easily genetically manipulated cell models of retinal cells. ARPE19 (ATCC CRL-2302) (Dunn et al., 1996), a spontaneously arising male retinal pigment epithelial cell line, and hTERT RPE-1 (ATCC CRL-4000) an hTERT immortalised female retinal pigment epithelial cell line are routinely used in molecular biology studies of retinal ciliopathies, owing to their retinal origin and ability to ciliate upon serum starvation and during late G1 (Spalluto et al., 2013). However, these cells are morphologically and functionally different from photoreceptors, and a dedicated photoreceptor cell line would be of enormous value in this area. Induced pluripotent stem cells (iPSCs) can be reliably differentiated into retinal photoreceptor cells following a 60-day differentiation protocol (Mellough et al., 2012). However, rapid loss of cells committed to a photoreceptor fate (CRX^+^/OPSIN^+^/RHODOPSIN^+^) is seen over days 45-60. The same phenomenon is observed in retinal progenitor cells from mice (Mansergh et al., 2010).

As an alternative to this, optic cups derived from mouse embryonic stem cells (Eiraku et al., 2011), human embryonic stem cells (Nakano et al., 2012) and human iPSCs (Meyer et al., 2011; Reichman et al., 2014) have become popular 3D models for retinal research. These take just 14-18 days to differentiate, and self-assemble in non-adherent culture. The resultant structures are homologous to embryonic retinal cups seen in vertebrate eye development, and include photoreceptors, but are not ideal models of mature retina. These also present problems associated with epigenetic effects, and the spheroid nature of optic cups which prevents access to the centre of these organoids for testing or analysis. Attempts to grow and differentiate ESCs and retinal progenitor cells into retinal sheets in custom matrices have been successful, but showed poor lamination (Worthington et al., 2016; Singh et al., 2017).

661W is an immortalised cone photoreceptor cell line derived from the retinal tumour of a mouse expressing SV40 T antigen (Tan et al., 2004). These cells have largely been used as a cell model for studying photo-oxidative stress and apoptosis, but not for studying inherited retinal dystrophies.

Here, we characterise the 661W cell line as a model for studying retinal ciliopathies.

## Materials and methods

### Cell culture

661W cells (Tan et al., 2004) were the kind gift of Prof Muayyad Al-Ubaidi, University of Houston. Cells were cultured in DMEM high glucose + 10% FCS at 37°C, 5% CO_2_, and split at a ratio of 1:5 once per week. hTERT-RPE1 cells (ATCC CRL-4000) were cultured in DMEM/F12 (50:50 mix) + 10% FCS at 37°C, 5% CO_2_, and split at a ratio of 1:8 once per week.

### Immunocytochemistry

Cells were seeded at 1 × 10^5^ per mL on sterile glass coverslips in complete media, and after 48 hours media was changed to serum-free media, and cells grown for a further 72 hours. Cells were rinsed in warm Dulbecco’s phosphate buffered saline (DPBS) and fixed in ice-cold methanol at −20°C for 5 minutes. Cells were then immediately washed with PBS, and incubated with blocking solution (1% w/v non-fat milk powder/PBS) for 15 minutes at room temperature. Coverslips were inverted onto primary antibodies in blocking solution in a humidity chamber and incubated at 4°C overnight. After 3 washes with PBS, cells were incubated with secondary antibodies and DAPI for 1 hour at room temperature in the dark. After 3 PBS washes and 1 dH_2_O wash, cells were mounted onto slides with Mowiol.

### Antibodies

#### Primary antibodies for IF

Mouse anti Polyglutamylated tubulin (GT335) 1:1000. Adipogen Life Sciences AG-20B-0020

Rabbit anti Gamma-tubulin 1:500. Abcam ab11317

Rabbit anti Arl13b 1:500. Proteintech 17711-1-AP

Rabbit anti Ift88 1:500. Proteintech 13967-1-AP

Rabbit anti Rpgrip1l 1:100. Proteintech 55160-1-AP

Mouse anti Cep164 1:100. Santa Cruz sc-515403

Rabbit anti long/medium wave pigment (green opsin) (Wang et al. 1992) 1:2000. Kind gift of Prof Jeremy Nathans, Howard Hughes Medical Centre, Johns Hopkins Medical School, Baltimore

Rabbit anti short wave pigment (blue opsin) (Wang et al. 1992) 1:2000. Kind gift of Jeremy Nathans, Howard Hughes Medical Centre, Johns Hopkins Medical School, Baltimore

#### Secondary antibodies for IF

Goat anti rabbit IgG AlexaFluor 488 1:1000

Goat anti mouse IgG AlexaFluor 568 1:1000

Donkey anti mouse IgG AlexaFluor 488 1:500

Donkey anti rabbit IgG AlexaFluor 568 1:500

#### Primary antibodies for WB

Mouse anti beta actin clone AC-15. 1:4000. Sigma-Aldrich A1978

Rabbit anti Ift88 1:500. Proteintech 13967-1-AP

Rabbit anti long/medium wave pigment (green opsin) (Wang et al. 1992) 1:5000. Kind gift of Prof Jeremy Nathans, Howard Hughes Medical Centre, Johns Hopkins Medical School, Baltimore

#### Secondary antibodies for WB

Donkey anti mouse 680 1:20,000 (LiCor)

Donkey anti rabbit 800 1:20,000 (LiCor)

Goat anti mouse HRP 1:5000 (Dako)

Goat anti rabbit HRP 1:5000 (Dako)

### High resolution confocal imaging

Confocal images were obtained at the Centre for Research in Biosciences Imaging Facility at UWE Bristol, using a HC PL APO 63x/1.40 oil objective CS2 lens on a Leica DMi8 inverted epifluorescence microscope, attached to a Leica SP8 AOBS laser scanning confocal microscope with 4 solid state AOTF supported lasers (405nm/50mW, 488nm/20mW, 552nm/20mW, 638nm/30mW), two standard PMTs and two high sensitivity HyD hybrid SMD GaAsP detectors. Images were captured using LASX software with Hyvolution II, with automated settings for highest resolution imaging, with pinhole set at 0.5AU. Images were deconvolved using Huygens Classic Maximum Likelihood Estimation (CMLE) algorithm (Scientific Volume Imaging). Images were assembled in Adobe Photoshop, and figures prepared using Adobe Illustrator.

### High-throughput confocal imaging

Images were obtained at Wolfson Bioimaging Facility University of Bristol, using the Perkin Elmer OperaLX high-throughput imager, using a 60x water objective lens, 405, 488 and 561nm lasers. PerkinElmer Opera Adjustment Plate containing multicolour beads was used to define the skewcrop and reference parameters. Cells were grown, fixed and immunostained in 96-well optical bottomed Cell Carrier plates (Perkin Elmer), and images were captured using OperaDB software. Individual z slices were exported as .flex files and assembled into maximum intensity projection z-stacks in Fiji ImageJ (Schindelin et al., 2012). These z-stacks were exported as 16-bit TIFFs, which were imported into CellProfiler for analysis using custom analysis protocols (Carpenter et al 2006; Kamentsky et al., 2011). Alternatively, z-stacks were constructed in OperaDB, and images analysed using PerkinElmer Acapella scripts, including algorithms ‘Find nuclei’, ‘Find cytoplasm’ and ‘Find spots’ (to find cilia).

### SDS-PAGE and western blotting

Total protein was extracted from cells using NP40 lysis buffer and scraping. Insoluble material was pelleted by centrifugation at 10,000 x g for 5 minutes and the total protein concentration of the supernatant was assayed using detergent compatible protein assay kit (BioRad). 20µg of total protein per sample was mixed with 2 x SDS loading buffer, boiled at 95°C for 5 minutes, and loaded onto pre-cast 4-12% NuPAGE Novex Bis-Tris gels (Life Technologies) alongside Spectra Multicolor Broad Range Protein ladder (Thermo Fisher). Samples were separated by electrophoreses in MES-SDS running buffer (Life Technologies) at 200V for 45 minutes. Protein was transferred to PVDF membrane using wet transfer at 40V for 2 hours. Membranes were incubated with 5% (w/v) non-fat milk/PBS to saturate non-specific binding, and incubated with primary antibody overnight at 4°C. After washing in PBST, membranes were incubated with secondary antibody for 1 hour at room temperature and exposed using 680nm and/or 780nm laser, or membrane was incubated with SuperSignal West Femto reagent (Pierce) and exposed using Chemiluminescence settings on LiCor Odyssey imaging system (LiCor).

### Scanning electron microscopy (SEM)

SEM images were obtained using a FEI Quanta FEG 650 field emission SEM at UWE Bristol Centre for Research in Biosciences Imaging Facility, using 20µs dwell, 30.00kV HV, 3.45µm HFW. 8.33e-7 Torr pressure, 60,000x magnification.

### RNA Sequencing

Total RNA was extracted from tissue using TRI reagent (Sigma Aldrich). RNA samples were treated with a TURBO DNA-free™ Kit (Ambion Inc.) using conditions recommended by the manufacturers, and then cleaned with an RNA Clean & Concentrator™-5 spin column (Zymo Research Corp.). RNA was tested for quality and yield using a NanoDrop 1000 spectrophotometer and an Agilent 2100 Bioanalyzer.

Six total RNA samples were supplied and prepared into sequencing libraries from ~500ng by Bristol Genomics Facility using the Illumina TruSeq Stranded mRNA kit. Briefly, RNA was polyA-selected, chemically fragmented to about 200 nt in size (4-minute fragmentation time), and cDNA synthesized using random hexamer primers. Each individual library received a unique Illumina barcode.

Quality of starting total RNA (diluted 1:100 to be within the limits of the assay) and the final libraries were also assessed using the Agilent TapeStation.

RNA-seq was performed on an Illumina NextSeq500 instrument with six libraries multiplexed and run across 4 lanes per flow-cell using 75 bp single end reads in high output mode. This resulted in more than 400 million reads per flow cell (651Mill, 590Mill PF), with an average of 94 million reads per sample.

Raw reads from 4 lanes per sample (4 FASTQ files) were aligned to the mouse (*Mus musculus*) full genome (GRCm38, UCSC mm10) using STAR, a splice-aware aligner (Dobin et al., 2013), with UCSC mm10.gtf gene model for splice junctions and the resultant BAM files were merged.

Again, using UCSC mm10.gtf file, raw gene counts were estimated on merged BAM files using HTSeq, using the union method and –stranded=reverse options (Anders et al. 2014). Differential gene expression was analysed using DESeq2 (Love et al., 2014) with statistical significance expressed as a p-value adjusted for a false discovery rate of 0.01 using Benjamini-Hochberg correction for multiple-testing.

GO Enrichment Analysis was carried out on the genes found to be differentially expressed between unstarved and starved cells using DAVID (Huang et al., 2009).

## Results

To confirm the cone photoreceptor origin of this cell line, and to fully characterise the expression profile of these cells, we performed whole transcriptome RNA sequencing on cells across 3 passages, with and without exposure to serum starvation (GEO, Supplementary Table 1). We show that these cells robustly express a range of markers of cone cell origin, including short wave (blue) and medium wave (green) opsin (Corbo et al., 2007), over a range of passages, in circumstances of both serum starvation and non-serum starvation (Figure 2a, Supplementary Table 2). Western blotting confirms expression of medium wave (green) opsin in the cytoplasmic fraction of these cells, with this band absent in RPE1 cells (Figure 2b). In keeping with the observation that photoreceptor gene expression falls on a continuous spectrum from cone-specific to rod-specific (Corbo et al 2007), we also saw low level expression of several rod-specific markers, including rhodopsin.

**Figure 2.**
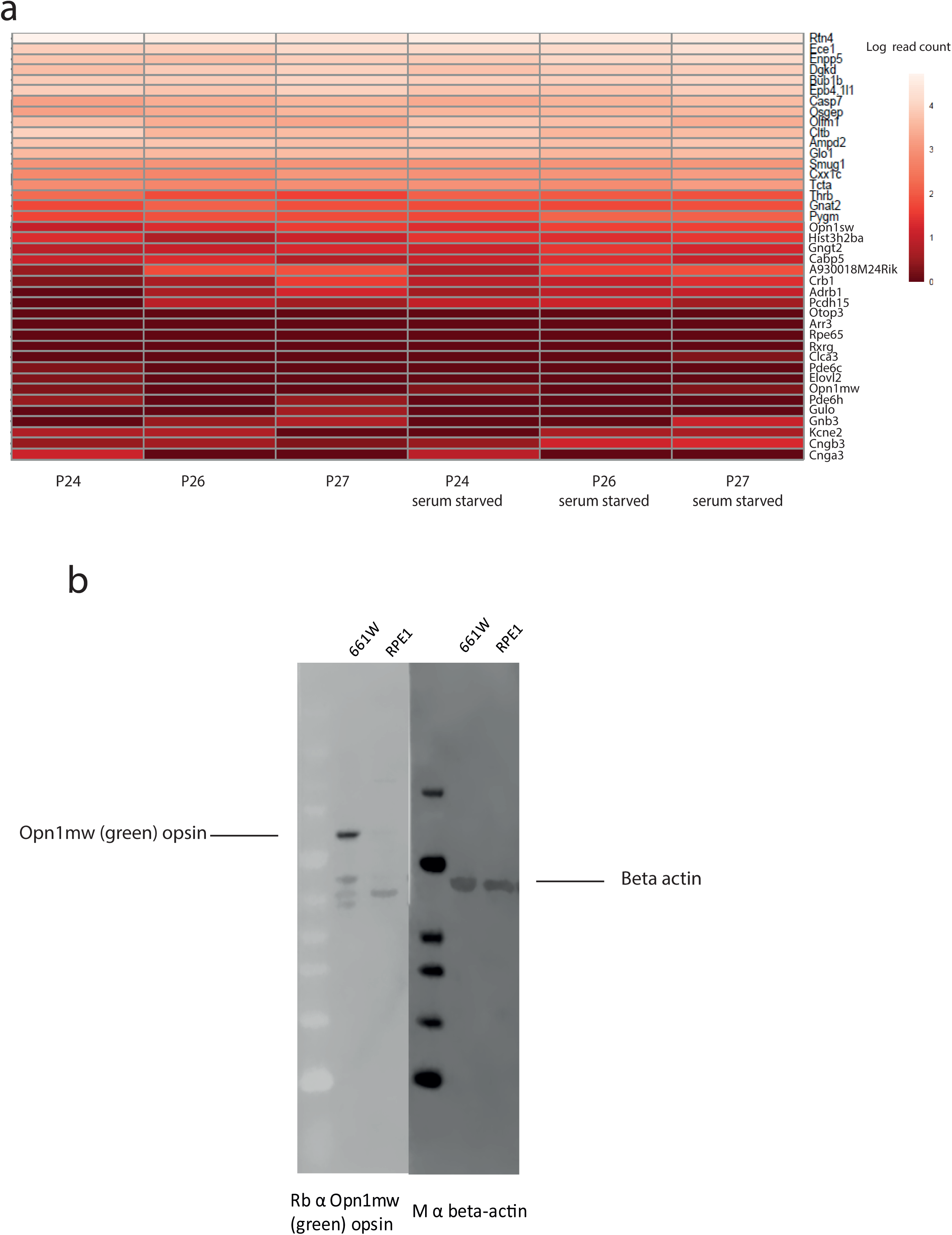
661W cells express markers of photoreceptor fate. (a) Heat map of expression of 40 genes characteristic of cone photoreceptors (Corbo et al., 2007) in 661W cells across 3 passages (P24, P26, P27) and in conditions of serum and serum starvation. Scale bar shows intensity of colour relative to log read count from HTSeq count outputs. (b) western blot showing specific expression of Opn1mw (green) opsin in 661W cytoplasmic extract, and absence of this band in RPE1 cells

In an attempt to differentiate these cells to develop a more photoreceptor-like morphology, we starved the cells of serum for 72 hours to induce cilia formation. Immunostaining and confocal microscopy of these cells shows that they grow long primary cilia, up to almost 15µm in length, and localise many cilium proteins to the axoneme (polyglutamylated tubulin, Ift88), membrane (Arl13b) and basal body (gamma tubulin) (Figure 3a-c). Whilst the axoneme of most primary cilia tends to be extensively post-translationally modified, in 661W cells only a small portion of the proximal axoneme is polyglutamylated (Figure 3a-c), consistent with a similar observation in cones *in vivo* (Eckmiller, 1996). Scanning electron microscopy (SEM) shows these cilia grow in characteristic ciliary pits in the cell membrane (Figure 3d). Furthermore, the cells localise both short wave and medium wave opsin to the base of this cilium (Figure 3e,f).

**Figure 3.**
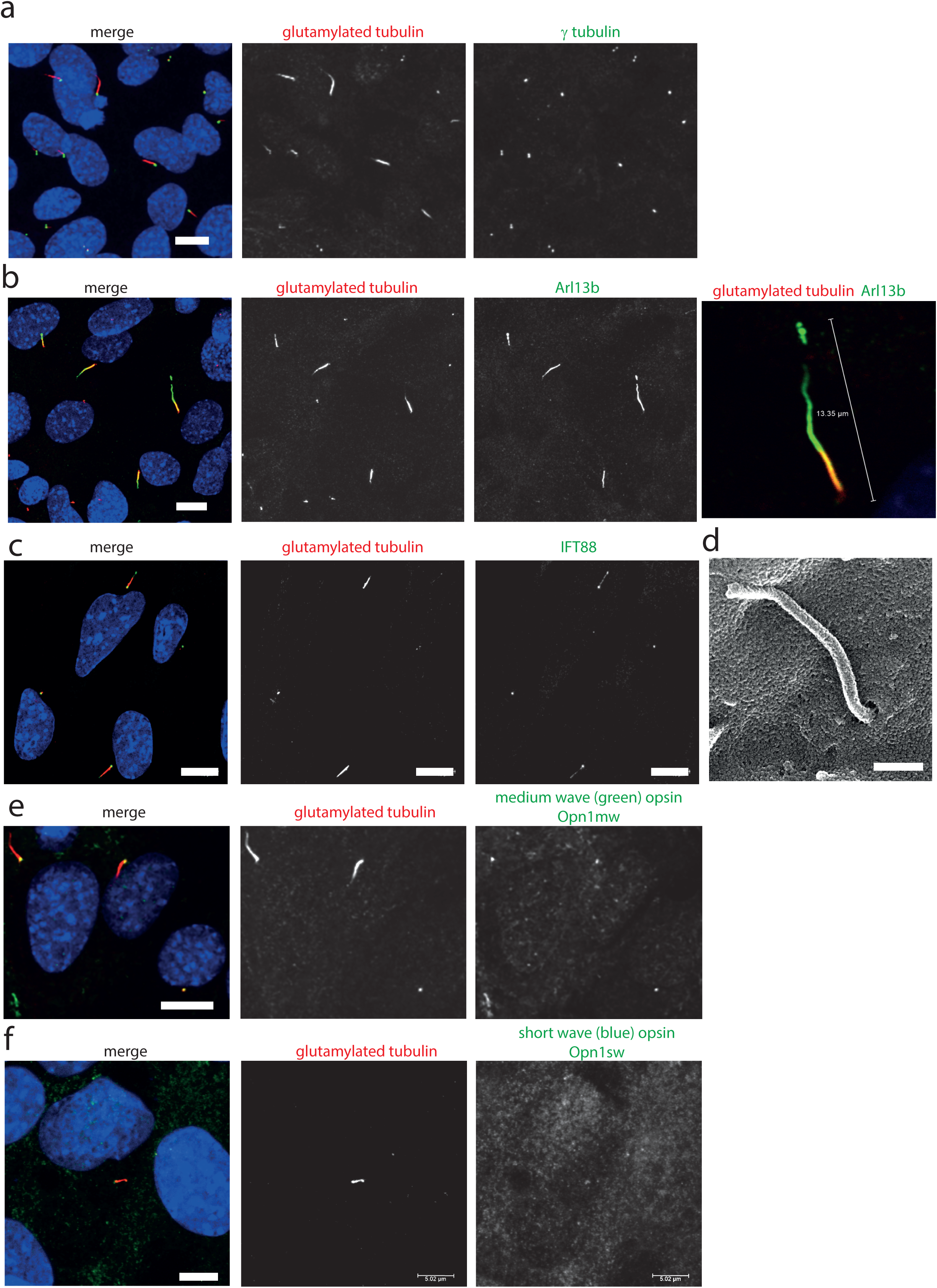
661W cells grow long primary cilia and localise opsins to the base of this cilium. (a) 661W stained with proximal axonemal marker polyglutamylated tubulin (red), basal body marker γ tubulin (green) show that many cells grow primary cilia. Cells counterstained with DAPI (blue). Scale bar 10µm. (b) Staining with cilium membrane marker Arl13b (green) shows that these cells grow cilia up to approx. 15µm long, with only the proximal portion of the axoneme polyglutamylated. Scale bar 10µm (c) Staining with IFT88 antibody (green) shows that this IFT protein localises along the cilium, with more concentrated localisation at the base and tip of the cilium. Scale bar 10µm. (d) Scanning electron microscope image of 661W cell, showing cilium in ciliary pit. Scale bar = 500nm. (e) Staining with Opn1mw and Opn1sw (f) show that these opsins localise to the base of the 661W cilium. Scale bars 5µm.

To further resolve the ultrastructure of this cilium, we used confocal microscopy with deconvolution to analyse the relative localisation of a centrosomal and basal body marker (gamma tubulin), distal centriolar appendage marker (Cep164), axonemal marker (Arl13b) and transition zone marker (Rpgrip1l) along the cilium (Figure 4a-c). This revealed an Arl13b cilium membrane extending from a ring of Cep164, lying distal to a ring of gamma tubulin of the mother centriole-derived basal body. A second gamma tubulin ring, of the daughter centriole, lies at approximately right-angles to this basal body. Rpgrip1l, a transition zone marker, extends from within the ring of Cep164 beyond this ring, presumably along the axoneme of the cilium. This differs from descriptions of Rpgrip1l localisation within the transition zone in primary cilia, where it is normally seen to reside in a tight ring narrower in diameter to Cep164, but more distal to Cep164 (Yang et al., 2015). However, it is known that transition zone assembly is highly cell-type specific, and Rpgrip1l plays a key role in this process (Wiegering et al., 2018). Extension of Rpgrip1l further beyond the distal appendages in this cell line is reminiscent of RPGR1P1L localisation in photoreceptors *in vivo*, where it is localised along the connecting cilium (Dharmat et al., 2018).

**Figure 4.**
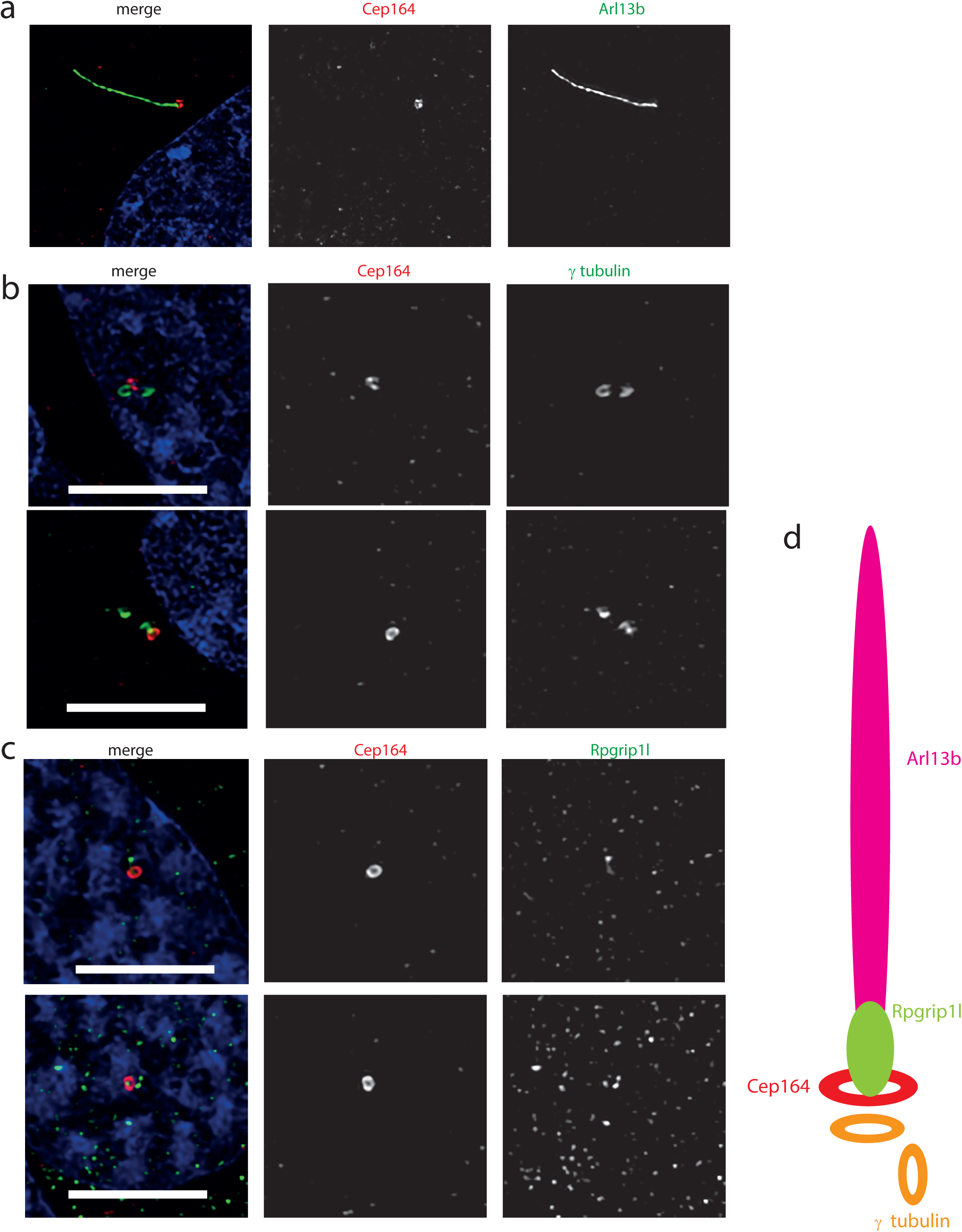
Ultrastructure of the 661W cilium. (a) Hyvolution imaging of distal centriolar appendage marker Cep164 (red), which localises specifically to the mature mother centriole of the basal body, and cilium membrane protein Arl13b (green). (b) Hyvolution imaging of distal centriolar appendage marker Cep164 (red), which localises specifically to the mature mother centriole of the basal body, and γ tubulin (green) which labels both mother and daughter centrioles. Scale bar = 5 µm. (c) Hyvolution imaging of distal centriolar appendage marker Cep164 (red), and transition zone protein Rpgrip1l (green). Scale bar = 5 µm. (d) Schematic representation of protein localisation in 661W cilia

To further investigate expression of cilia genes in this cell line, we extracted HTSeq count data from our RNAseq data for Syscilia Gold Standard (SCGS) genes (van Dam et al., 2015) in each replicate of starved and unstarved cells. The cells showed robust expression of 281 SCGS genes in unstarved and starved cells across all replicates, and statistically significant (p<0.05) upregulation of 38/281 (13.5%) of these genes upon serum starvation (Supplementary Table 3). This includes 12 genes known to be causative of retinal ciliopathies (Figure 5a). Serum starvation also induced statistically significant upregulation of several cilia genes linked to ciliopathies which do not have a retinal phenotype (Figure 5b).

**Figure 5.**
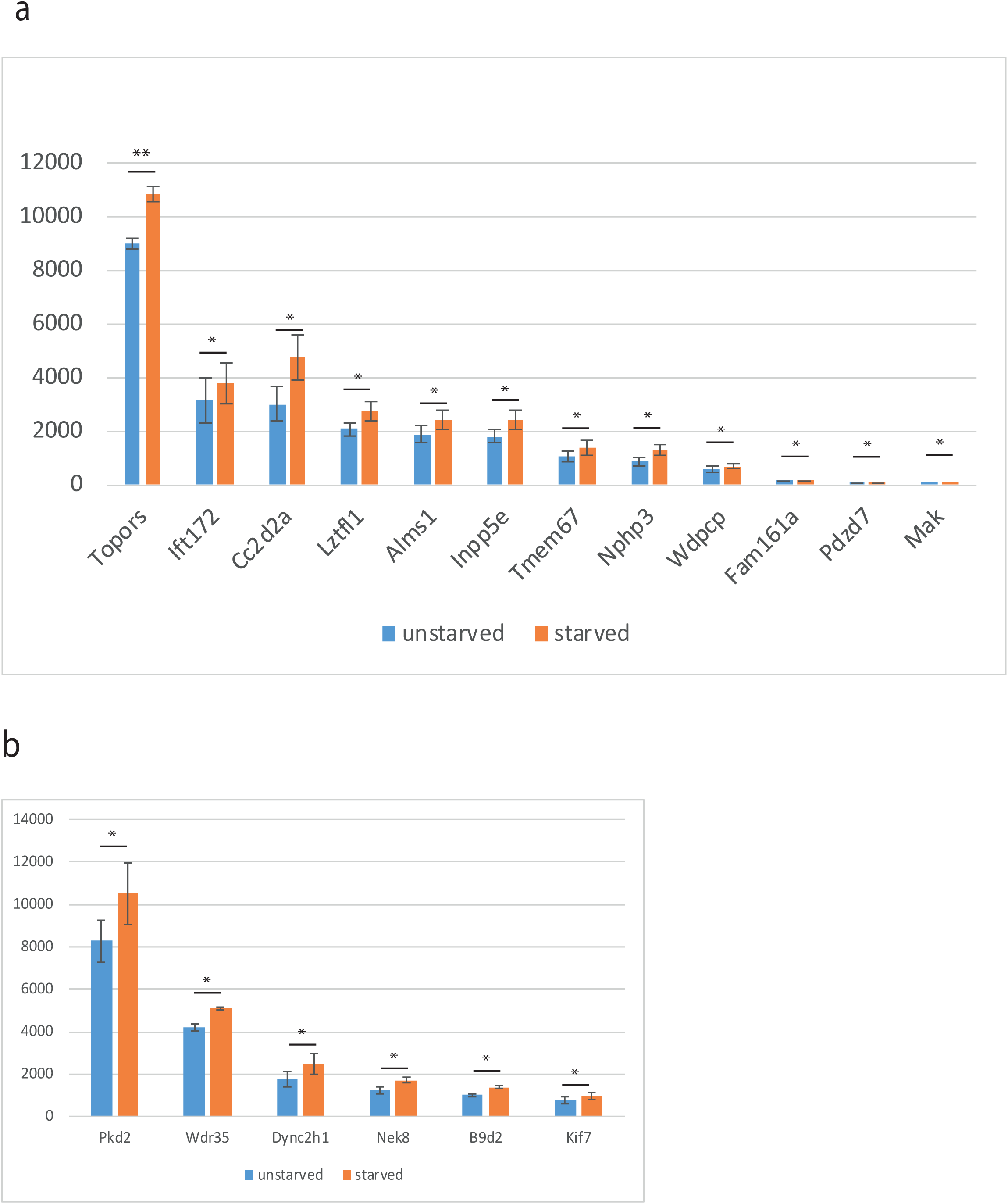
Expression of a subset of Syscilia Gold Standard (SCGS) genes mutated in retinal ciliopathy patients is upregulated upon serum starvation of 661W cells. (a) Graph showing mean HTSeq read count of selected Syscilia Gold Standard (SCGS) genes mutated in retinal ciliopathies, in unstarved and serum starved cells, to show upregulation of these genes upon serum starvation to induce cilium formation. ** p<0.005 * p<0.05. (a) Graph showing mean HTSeq read count of selected Syscilia Gold Standard (SCGS) genes mutated in other ciliopathies, in unstarved and serum starved cells, to show upregulation of these genes upon serum starvation to induce cilium formation. * p<0.05.

We then used DESeq2 to identify all genes differentially expressed in different conditions (starved vs unstarved), controlling for batch-effects, and performed enrichment analysis on all hits with an adjusted p-value of >0.01 using DAVID (Huang et al., 2009) (Supplementary Table 4). This identified a number of enriched annotation clusters, including those with the Gene Ontology (GO) terms ‘neuron differentiation’ (enrichment score 2.22), ‘eye development’ (enrichment score 0.16) and ‘sensory perception’ (enrichment score 0.52), ‘GPCR, rhodopsin-like superfamily’ (enrichment score 0.25) and ‘centrosome’ (enrichment score 0.010) (Supplementary Table 5).

Non-coding RNA data from RNA sequencing of this cell line identifies a number of novel RNAs not previously linked to photoreceptor fate (Supplementary Table 6). Most noteworthy is mmu-mir-6236, which was expressed at very high levels in starved and unstarved cells. This miRNA has not been previously linked to photoreceptor fate or function. These cells were also found to express miR-17-92, which has been previously linked to neuronal differentiation (Bian et al., 2013; Mao et al., 2016), and may contribute to neuronal photoreceptor differentiation in these cells. Interestingly, expression of this microRNA has been shown to promote cyst growth in polycystic kidney disease, a common ciliopathy (Patel et al., 2013). It could be hypothesised that this miRNA plays a role in ciliogenesis in both the kidney and the photoreceptor. The only non-coding RNA to show a more than 2-fold increase in expression upon serum starvation was miRlet7b, a highly abundant regulator of neuronal differentiation in the brain (Lagos-Quintana et al 2002; Berezikov et al., 2006; Roush and Slack, 2008).

Finally, in order to evaluate the utility of 661W cells for high-throughput screening, we performed siRNA knockdown of cilia protein Ift88 in 661W cells in 96 well optical-bottom culture plates, and assayed cell number and cilia number by high-content imaging, using Arl13b and polygutamylated tubulin as cilia markers (Figure 6a). Nuclei were identified using DAPI, and cilia were detected using a modified ‘find spots’ Perkin Elmer image analysis algorithm. This showed that knockdown of positive control Plk1 induced a statistically significant reduction in cell number (Figure 6b), and Ift88 knockdown induced a statistically significant reduction in the number of cells with a single cilium (Figure 6b,c).

**Figure 6.**
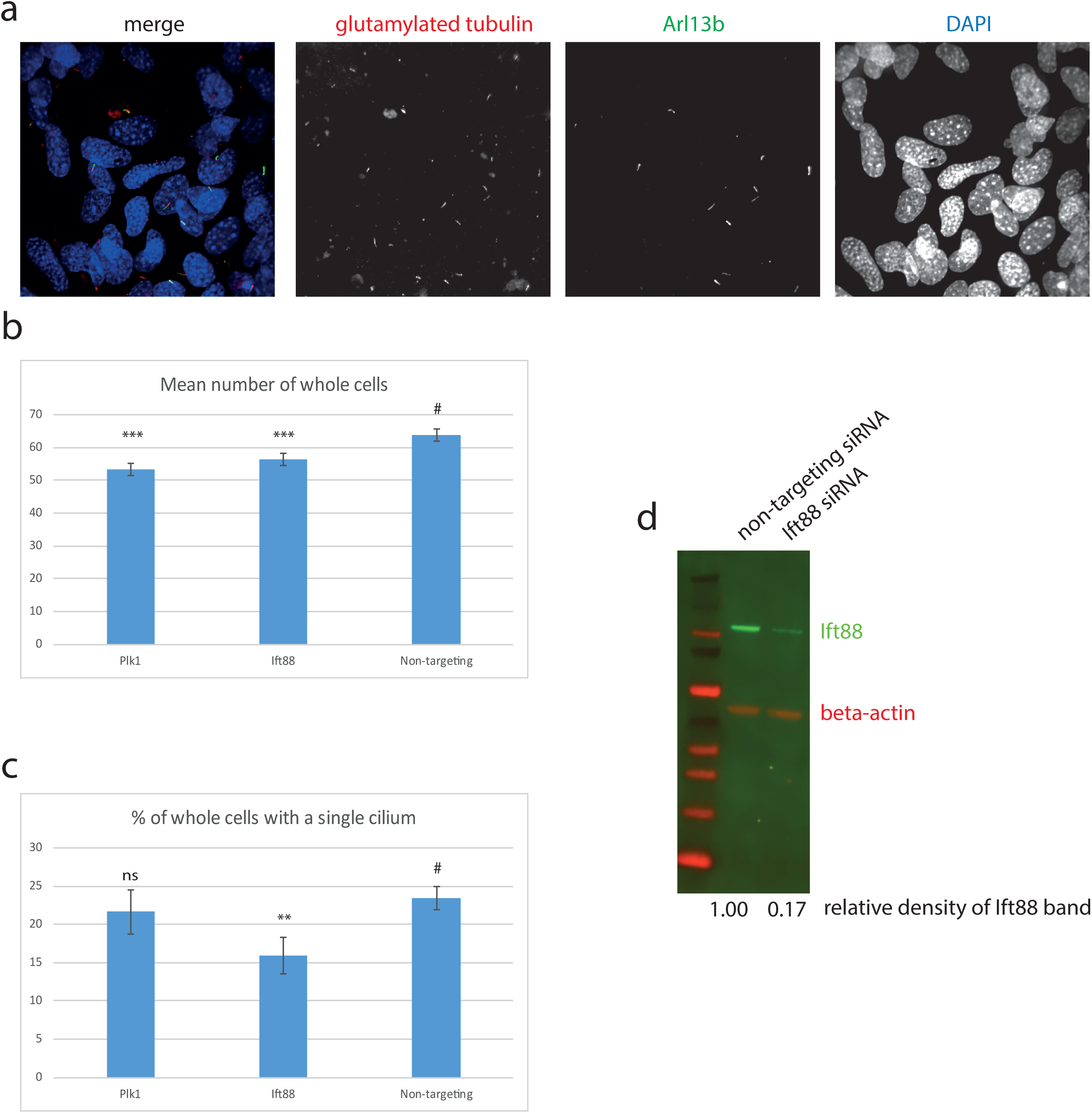
High content imaging of 661W cells. (a) Example images from the Perkin Elmer Opera, showing 661W cells stained wth glutamylated tubulin (red), Arl13b (green) and DAPI (blue). (b) Mean number of whole cells per field of view is statistically significantly reduced after 72 hr Plk1 and Ift88 siRNA knockdown compared to cells treated with non-targeting control. ***p <0.0001 (c) % of cells with a single cilium per field of view is statistically significantly reduced after 72 hr Ift88 siRNA knockdown compared to cells treated with non-targeting control. **p <0.001. (d) western blotting confirms reduction in Ift88 protein level (green) after 72hr Ift88 siRNA knockdown, compared to non-targeting control. Red shows beta actin loading control.

## Discussion

Our comprehensive deep sequencing of total mRNA and selected non-coding RNAs in the 661W cell line confirms its cone photoreceptor origin, and ability to grow a photoreceptor-like cilium in culture after serum starvation (Figure 2, 3, 4, Supplemtary Table 1). A recent paper described the 661W cell line as a retinal ganglion-like cell line, owing to its expression of markers specific to retinal ganglion cells, such as Rbpms, Pouf2, Pouf3, Thy1 and γ-synuclein (Sayyad et al., 2017). Our whole transcriptome RNA sequence data confirms that this cell line expresses Rbpms, Thy1 and Sncg at low levels, but does not express Pouf2 or Pouf3 (Supplementary Table 1). This suggests that this cell line does indeed have some features of retinal ganglion cells, but overwhelmingly expresses markers of cone photoreceptor fate. Our data supports the conclusion of Sayyed and colleagues, that this cell line shows properties of both retinal ganglion and photoreceptor cells, and is a useful *in vitro* photoreceptor model.

Immunofluorescence confocal microscopy and deconvolution of images of these cells reveals a cilium similar in structure to cone photoreceptor cilia *in vivo*. The cell extends a very long (~15µm) cilium, which is only post-translationally modified at the proximal end, and localises Rpgrip1l along this proximal region, and opsins to the base of the cilium (Figure 3, 4). The cell line expresses many bona fide cilia genes, including many mutated in retinal ciliopathies, and expression of many of these increases upon serum starvation of the cells (Figure 5, Supplementary Table 3, 4). Knockdown of key cilia gene Ift88 results in robust reduction in percentage of whole cells with a single cilium, which can be readily assayed by high-content imaging (Figure 6).

Our demonstration of high-content imaging of 661W cells illustrates the potential utility of this cell line for high-throughput screening. A recent high-throughput small molecule screen in RPE1 cells successfully identified eupatilin as a small molecule which rescued transition zone defects in CEP290 knockout RPE1 cells (Kim et al., 2018). A similar approach using 661W cells could be an even more clinically relevant method for identifying small molecules which could be used for treating retinal ciliopathies. Similarly, the cell line could be useful for reverse genetics functional genomics screening to identify novel retinal cilia genes and retinal ciliopathy candidate genes, similar to previous screens (Wheway et al., 2015). These methods are of enormous importance in a field where effective treatments are so limited.

In summary, we provide evidence that 661W cells are a useful *in vitro* model for studying retinal ciliopathies.

## Supporting information

## Author contributions

GW conceived of the study, performed experiments, carried out data analysis, imaging, image analysis, bioinformatics analysis, and wrote the paper. LN performed experiments, carried out data analysis, imaging and image analysis. DN contributed to bioinformatics analysis. SC provided support in high-throughput imaging and image analysis.

## Acknowledgements

We would like to thank Dr David Patton for assistance with SEM.

We would like to thank Prof Muayyad Al-Ubaidi for the gift of the 661W cells, and Prof Jeremy Nathans for the gift of the opsin antibodies.

We would like to thank all at Scientific Volume Imaging for assistance with Huygens deconvolution

GW would like to thank Prof David Stephens and Prof Catherine Nobes for providing access to the laboratories at University of Bristol School of Biochemistry.

